# Modeling intrahippocampal effects of anterior hippocampal hyperactivity relevant to schizophrenia using chemogenetic excitation of long axis-projecting mossy cells in the mouse dentate gyrus

**DOI:** 10.1101/2020.12.15.422938

**Authors:** James P. Bauer, Sarah L. Rader, Max E. Joffe, Wooseok Kwon, Juliana Quay, Leann Seanez, Chengwen Zhou, P. Jeffrey Conn, Alan S. Lewis

**Author notes:** **Corresponding author:** Alan S. Lewis, MD, PhD, Department of Psychiatry and Behavioral Sciences, Vanderbilt University Medical Center, 465 21st Ave S., MRBIII Room 6140B, Nashville, TN 37240.

## Abstract

**Background:** The anterior hippocampus of individuals with early psychosis or schizophrenia is hyperactive, as is the ventral hippocampus in rodent models for schizophrenia risk. Hyperactive ventral hippocampal projections to extrahippocampal brain regions contribute to schizophrenia symptoms, but less is known about the functional effects of hyperactive projections within the hippocampal formation long axis. We approached this question by testing whether hyperactivation of ventral dentate gyrus (DG) mossy cells (MCs), which densely project intrahippocampally to the dorsal DG, influences spatial memory, a cognition dependent on intact dorsal DG function.

**Methods:** In CD-1 mice, we targeted dorsal DG-projecting ventral DG MCs using an adeno-associated virus intersectional strategy. In vivo fiber photometry recording of ventral DG MCs was performed during exploratory behaviors. We targeted excitatory chemogenetic constructs to ventral DG MCs and tested whether their hyperactivation impaired encoding in a spatial memory task.

**Results:** Ventral DG MCs were activated during behavior related to environmental information gathering (rearing) but not during non-exploratory motor behaviors. Ventral DG MCs made functional monosynaptic inputs to dorsal DG granule cells, with chemogenetic activation of ventral DG MCs leading to increased activity of dorsal DG granule cells. Finally, chemogenetic activation of ventral DG MCs during the encoding phase of an object location memory task impaired retrieval 24 hours later, without effects on locomotion or other exploratory behaviors.

**Conclusions:** These data suggest that localized hippocampal hyperactivity may have longitudinal intrahippocampal functional consequences, supporting study of longitudinal circuits as targets to mitigate cognitive deficits associated with schizophrenia.

## INTRODUCTION

Cognitive impairment in schizophrenia (CIIS) independently predicts functional deficits (1-3). However, there are currently no FDA-approved pharmacotherapies for CIIS, suggesting underlying mechanisms are incompletely understood. Here we aim to harness the power of molecular tools to manipulate neural circuits in rodents to identify underlying mechanisms accounting for robust neuroimaging findings in patients with schizophrenia or early psychosis. Several independent groups have reported that the anterior hippocampus in such patients is reduced in volume (4-8) and hyperactive at baseline (4, 9-12). It was recently shown in early psychosis patients that greater baseline hyperactivity was associated with reduced anterior hippocampal recruitment during cognitive tasks (9). However, the exact relationship between aberrant anterior hippocampal activity and deficits in learning and memory remains unclear (13).

Many rodent models with genetic or developmental insults for modeling schizophrenia etiology also exhibit hyperactivity in the ventral hippocampus (vHPC), the rodent analogue of the human anterior hippocampus (14, 15). Previous work in these models has shown that vHPC hyperactivity drives aberrant dopamine neurotransmission through extrahippocampal projections to nucleus accumbens (reviewed in (16)) and disrupts distinct cognitive domains, such as spatial working memory, via projections to medial prefrontal cortex (17, 18). Much less is known about how hyperactivity in the anterior portion of the HPC might directly interact with the posterior HPC (analogous to the rodent dorsal HPC (dHPC)) via HPC long axis circuitry. This question is important because certain cognitive deficits present in individuals with schizophrenia are dependent on posterior HPC function, such as visuospatial contextual memory (19-22). Using functional techniques in mice may identify the cognitive consequences of hyperactivity of vHPC intrahippocampal circuits, similar to studies identifying extrahippocampal ramifications.

To model hyperactive long-distance projections originating in the ventral half of the HPC, here we focus on ventral mossy cells (vMCs), which are glutamatergic neurons with cell bodies in the dentate gyrus (DG) hilus (23, 24). Ventral MCs send long-distance excitatory projections to both ipsi- and contralateral DG, resulting in dense connectivity of the entire longitudinal hippocampal axis (25-28). Inhibition of vMCs impairs spatial memory encoding (29), a surprising finding given the reliance on the dHPC for spatial memory (30-34). Mossy cells are sensitive to even low levels of activity throughout the HPC (35), and thus are plausible candidates to be functionally responsive to HPC hyperactivity. In this study, we used vMCs as a longitudinal HPC model circuit by examining their in vivo activity during exploratory behavior, testing their connectivity with target dDG granule cells, and determining the consequences of chemogenetic vMC hyperactivation in a spatial memory task that is dependent on dHPC function.

## MATERIALS AND METHODS

Detailed methods are reported in the Supplemental Information.

### Animals

Wildtype CD-1 male and female mice (Charles River Laboratories, Wilmington, MA; age at delivery: six-eight weeks) were acclimated to the vivarium for at least one week. All mice were group housed except for cannula-implanted mice, which were single housed. Wildtype C57BL/6J male mice (Jackson Laboratory, Bar Harbor, ME; age at delivery: six-seven weeks) were used as intruders during resident-intruder social interactions. Mice were housed under standard conditions with a 12 h light–dark cycle (lights on at 6:00 am) with food and water available ad libitum. The Vanderbilt Institutional Animal Care and Use Committee approved all procedures.

### Drugs

Clozapine N-oxide (CNO) for systemic administration was provided by the NIMH Chemical Synthesis and Drug Supply Program, dissolved in 0.5% dimethyl sulfoxide in saline, and injected intraperitoneally (i.p.) 30 mins prior to start of behavioral assay. For slice electrophysiology, CNO was purchased from Hello Bio (Princeton, NJ).

### Adeno-associated viruses

Adeno-associated virus particles (AAVs) used, purchased from Addgene (Watertown, MA), were AAV.pgk.Cre, pAAV.Syn.Flex.GCaMP6f.WPRE.SV40 (36), pAAV-hSyn-DIO-mCherry, pAAV-hSyn-DIO-hM3D(Gq) (37), pENN.AAV.hSyn.Cre.WPRE.hGH, and pAAV-EF1a-double floxed-hChR2(H134R)-mCherry-WPRE-HGHpA.

### Stereotaxic surgery and viral infusion

Under inhaled isoflurane anesthesia, mice were mounted in a stereotaxic frame, and AAV (0.2 – 0.4 μL depending on brain region) was infused at a rate of 50 nL/min using a 2 μL syringe (Hamilton, Reno, NV). Dorsal DG: anterior/posterior (AP): −1.94 mm, medial/lateral (ML): 1.20 mm left; dorsal/ventral (DV): −2.50 mm. Ventral DG hilus: AP: −3.40 mm, ML: ±3.00 mm, DV: −3.50 mm. Fiber photometry cannulas were targeted to the same coordinates as ventral DG hilus and cemented in place with C&B-Metabond (Parkell, Edgewood, NY). Mice were housed for at least three weeks prior to experimentation to allow for viral expression. Targeting was confirmed by fluorescence microscopy. Mice with either unilateral or bilateral vMC expression were included in behavioral analyses due to extensive bilateral projections.

### Fluorescent immunostaining, microscopy, and cell counting

Primary antibodies for 40 μm free-floating sections were mouse anti-calretinin (MAB1568, Millipore, Burlington, MA (38)) and rabbit anti-cFos (226 003, Synaptic Systems, Goettingen, Germany (39)), both diluted 1:1000. Widefield fluorescent images were acquired with an AF6000 LX fluorescent microscopy system (Leica, Buffalo Grove, IL) equipped with an HCX PL FLUOTAR 10x objective (NA = 0.30). Confocal images were acquired with an LSM 710 META Inverted microscope (Zeiss, White Plains, NY) equipped with a 20x Plan-Apochromat objective (NA = 0.8). Cell counting was performed blinded from confocal images.

### Electrophysiology

To validate DREADD activation of vMCs, mice were rapidly decapitated under deep isoflurane anesthesia and 300-µm horizontal sections containing the vDG were prepared in *N*-methyl-D-glucamine cutting solution and recorded in artificial cerebrospinal fluid. Hilar MCs were identified by mCherry expression, patched to obtain whole cell access, and voltage-clamped at −75 mV while 10 µM CNO was perfused in the bath. The mean of the holding current was quantified before and after CNO perfusion using pClamp 10.4 software (Axon Instruments, Union City, CA). For optogenetic activation of vMC terminals in dDG, brain slices were prepared as previously described (40, 41). Whole-cell patch-clamp recordings were made from dDG granule cells and slices were continuously superfused with ACSF. Current-clamp mode was used to record action potentials. 473 nm blue laser light (5 ms pulse duration, 20 Hz, 1 s duration) was delivered through a fiberoptic cable to stimulate channelrhodopsin-expressing vMC axon terminals. In both sets of experiments, recordings were acquired with a Multiclamp 700B amplifier (Molecular Devices, Sunnyvale, CA), filtered at 2 kHz and digitized at 10 kHz.

### Behavioral studies

#### General

Mice were habituated to the dimly lit testing room for at least one hr before testing. Testing occurred between 0900 and 1700. Experimental and control groups were tested on the same days and all tests were videotaped. Vehicle or CNO was injected i.p. 30 mins prior to the start of behavioral assays.

#### Open field test

Mice were placed into the center of a 61 cm by 61 cm square arena for 10 mins. Distance traveled, time spent in arena center, and number of center entries were quantified using ANY-maze version 6.1 (Stoelting, Wood Dale, IL). Rearing, defined as standing upright on hind legs, was quantified manually by an observer blind to treatment group.

#### Object location memory test

Performed largely as described previously (42). Mice were handled for about two mins daily for five days, followed by five mins of habituation daily in an empty 61 cm by 61 cm open field containing spatial cues on the wall for six days (final two days of handling were combined with habituation). The next day, mice underwent an encoding session during which they were placed in the same arena now containing two identical 500 mL glass bottles spaced ∼30 cm apart and allowed to explore the arena for 10 mins. Bottles were cleaned with dilute alcohol solution between mice. 24 hrs later, mice were returned to the same arena for a 5 min retrieval session where one bottle remained in the same location and the other bottle was moved ∼40 cm to a novel location. The side from which the bottle was moved was counterbalanced across mice. Time spent exploring each bottle was quantified from video by blinded observers. Only time during which mice directly explored the object (∼0-2 cm away) and were not rearing on hindlegs was counted (42). Discrimination index (DI) was quantified as (time spent exploring novel object – familiar object)/(total time spent exploring both objects). Mice demonstrating an initial side preference during the encoding session (DI > 0.20) (29) were removed from the analysis (n = 2 mCherry, 3 hM3Dq).

#### cFos expression

Mice were habituated to the testing room for >2 hours, administered CNO or vehicle i.p., and transcardially perfused 90 minutes later.

#### Resident-intruder test for fiber photometry

Performed similarly to that previously described (43). To start the assay, a C57BL/6J mouse was introduced to the home cage of the photometry cannula-implanted single housed CD-1 male and allowed to interact for >10 mins.

### Fiber photometry

Photometry was performed from male CD-1 mice implanted with photometry cannulas (Doric Lenses, Quebec, Canada) using Doric Lenses system controlled by Doric Neuroscience Studio (DNS) version 5.3. Video recording was time-locked with photometry signal. Signal processing was performed by DNS photometry analyzer. ΔF/F_0_ was calculated for the 405 nm and 465 nm channels independently using a least mean square fit, and then ΔF/F_0_ (405 nm) subtracted from ΔF/F_0_ (465 nm) to yield a corrected trace that was subsequently lowpass filtered at 2 Hz (44) to yield the final bulk calcium signal. This signal was Z-normalized across the entire social interaction. Behavior in videos was manually annotated for exploratory rearing (centered around apogee) and offensive attack onset. Data analysis was performed using custom written scripts in Matlab R2019a (MathWorks, Natick, MA).

### Statistical analysis

To compare two groups, unpaired t tests with Welch’s correction or paired t tests were used, as appropriate. To compare three or more groups, one-or two-way ANOVA with Sidak’s multiple comparisons test was used. One-sample t tests were used to compare experimental to hypothetical means. p < 0.05 was considered significant and all tests were two-tailed. Statistical analyses were performed in Prism 9 (Graphpad, San Diego, CA). Error bars depict standard error of the mean (s.e.m.).

## RESULTS

### Targeting vMCs projecting to the dDG

We exploited the well-organized distant bilateral axonal projection of vMCs to the inner molecular layer of the dDG (**Figure 1A**) (25, 27, 45-47) to express proteins of interest in longitudinal-projecting vMCs in wildtype mice. Retrograde-AAV-pgk-Cre was unilaterally infused into dDG and Cre-dependent GCaMP6f was infused into the contralateral vDG hilus (**Figure 1B**). This resulted in the expression of GCaMP6f in large ventral hilar cells, with a band of GCaMP6f+ projections to the contralateral vDG and bilateral dDG inner molecular layers (**Figure 1C**). Our stereotaxic targeting of vMCs was selected to be consistent with previous functional studies of both vMCs (29) and vDG granule cells (48), and conforms with a recent detailed anatomical study of MC projections which defined the dDG as the rostral half of the DG (AP −1.0 to −2.5 mm) and the vDG as the caudal half of the DG (AP −2.7 to −4.0 mm) (49). Dorsal and vMCs differ in their expression of calretinin, which in mice is only expressed in vMCs (**Supplementary Figure 1**) (27, 28, 50). Confirming that our targeting coordinates were consistent with previous definitions of vMCs, in the hilus we found ∼87% of GCaMP6f+ neurons were also calretinin+, while ∼68% of calretinin+ neurons were GCaMP6f+. We did identify low levels of GCaMP6f off-target expression in neighboring vDG granule cells and vCA3 pyramidal neurons (n = 4 mice; **Figure 1D, E**). Thus, these data show that this strategy is selective but not perfectly specific for vMCs, similar to other methods for MC targeting using D2-Cre (51) or Crlr-Cre (52) transgenic lines (52, 53).

**Figure 1.**
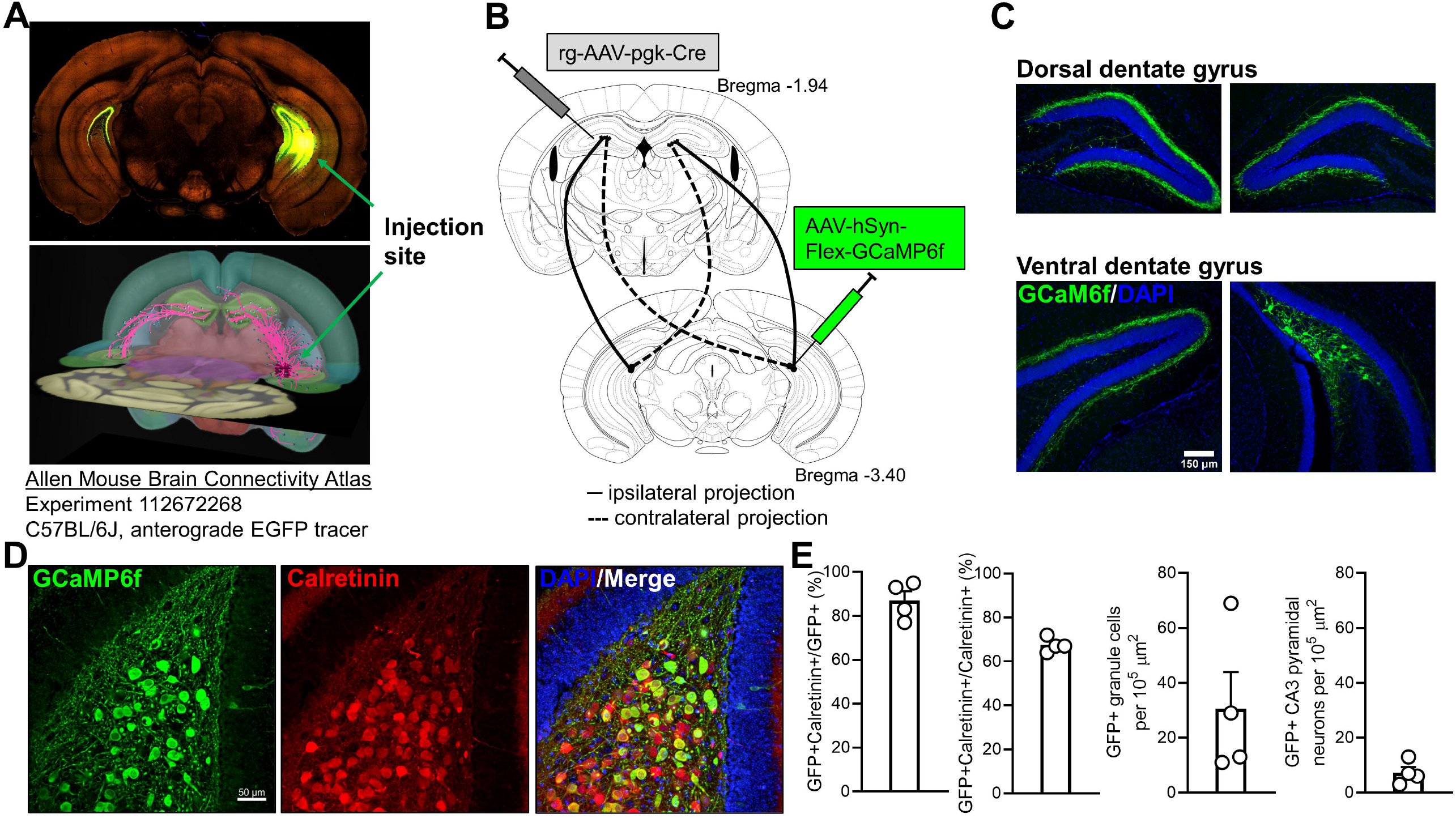
Intersectional targeting of longitudinal ventral mossy cell projections. A Ventral mossy cell (vMC) to dorsal dentate gyrus (dDG) circuit of interest as illustrated by the Allen Mouse Brain Connectivity Atlas (88). AAV expressing EGFP was infused into ventral hilus of wildtype C57BL/6J mouse (top). Serial two-photon tomography shows axonal projections throughout longitudinal extent of hippocampus, with strong projection to dDG inner molecular layer (https://connectivity.brain-map.org/projection/experiment/112672268). **B** To target vMCs, retrograde-AAV-pgk-Cre was infused into dDG inner molecular layer, and AAV-hSyn-Flex-GCaMP6f was infused into contralateral vDG hilus, enabling recombination only in vMCs that project to dDG. **C** Fluorescence microscopy from CD-1 mouse following targeting strategy shown in (B) reveals GCaMP6f is expressed in vMC somata with organized projections to contralateral vDG and bilateral dDG. **D** GCaMP6f from targeting strategy in (B) is highly colocalized with calretinin, a marker for ventral but not dorsal MCs. **E** Quantification of viral targeting strategy in (B) from n = 4 mice reveals most GCaMP6f+ hilar neurons are calretinin+, with limited off-target expression in nearby ventral granule cells and CA3 pyramidal neurons.

### Activation of longitudinally-projecting vMCs during behaviors related to environmental information gathering

The dDG is a critical brain region for spatial information processing in the rodent (54). While recent in vivo studies have shown dMCs play key roles in spatial coding (55-57), less is known about the in vivo activity of vMCs. Because of the longitudinal connectivity between vMCs and the dDG, we wondered whether vMCs might also contribute environment-based input to the dDG. We used fiber photometry (58, 59) to record bulk GCaMP6f calcium activity in vMCs (**Figure 2A**) as a proxy for neuronal firing during exploratory rearing events. Rearing on hind legs rapidly changes the mouse’s vantage point and multisensory inputs from local cues to more global environmental landmarks thereby representing a period of environmental information gathering (60, 61) and is associated with marked changes in DG activity, including in the hilus (62). We found that vMC activity increased concurrent with rearing apogee (**Figure 2B**), which was consistent within (**Figure 2C**, 46 rearing events in one mouse) and across (**Figure 2D**, 158 rearing events from 5 mice) mice. This activity increase might simply result from the abrupt onset of movement inherent in rearing events. To test whether the abrupt onset of movement in another behavior that is not exploratory in nature yielded the same activity increase, we examined vMC activity during the onset of offensive attack during social interaction. Despite offensive attack involving rigorous onset of movement, we found no consistent change in vMC activity associated with attack behavior in the same mice during the same behavioral session (**Figure 2E**, 50 attacks from 5 mice). Ventral MC activity was significantly greater during rearing compared to attack (t(4) = 4.02, p = 0.016) (**Figure 2F**). These experiments suggest that vMCs projecting to the dDG are activated during environmental information gathering, but not non-specifically activated by any locomotor behavior.

**Figure 2.**
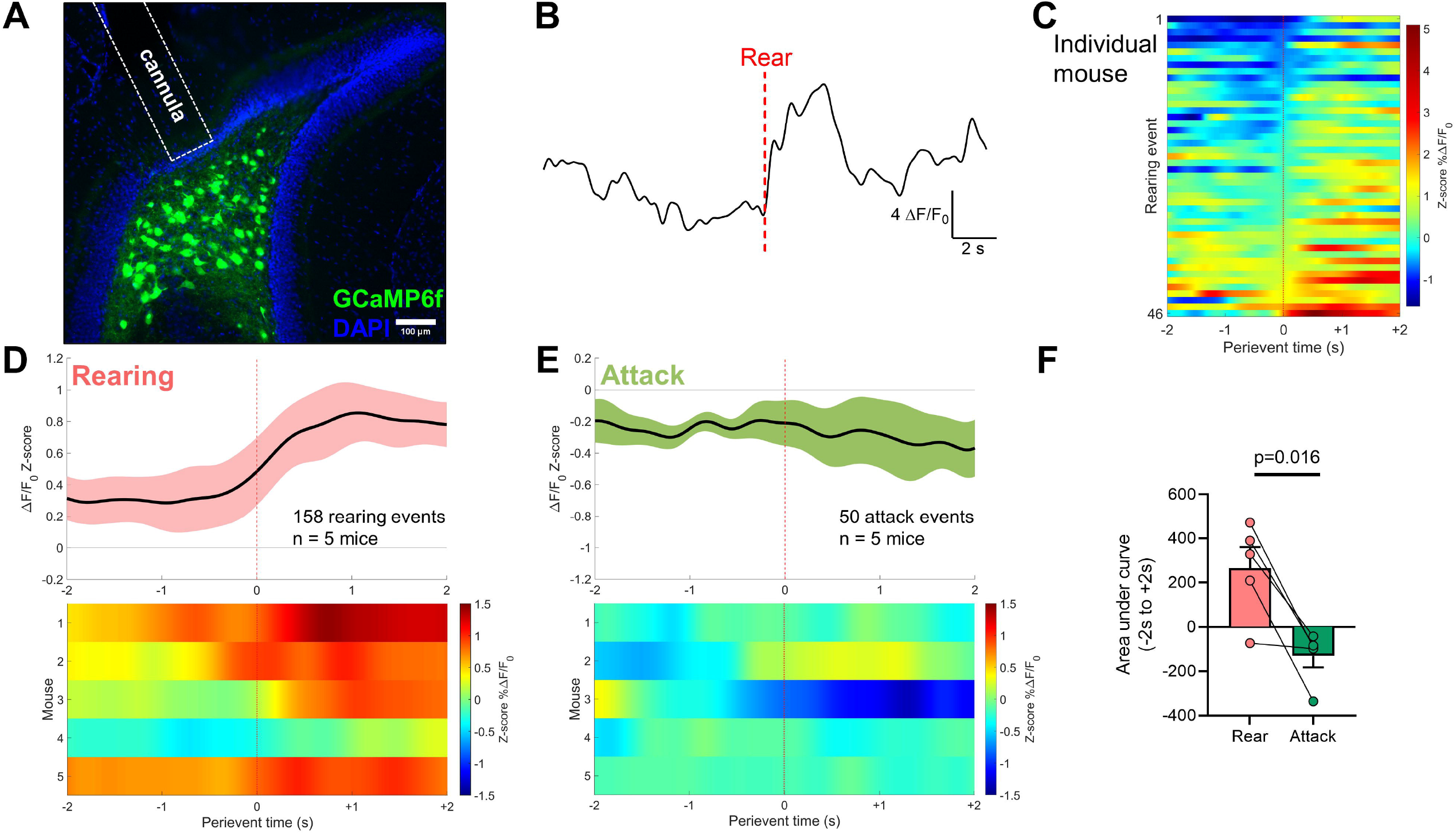
In vivo fiber photometry reveals ventral mossy cells are selectively activated by behavior related to environmental information gathering. A Fluorescence microscopy of GCaMP6f expression in ventral mossy cells (vMCs) using the intersectional targeting strategy and fiber optic cannula placement. **B** Individual photometry trace surrounding an example exploratory rearing event. **C** ΔF/F_0_ Z-scores of rearing events aligned to time of rearing apogee in an individual mouse. Events are ordered by Z-score and consistently show increased photometry signal during rearing. **D** ΔF/F_0_ Z-scores for N = 5 mice were aligned to time of rearing apogee, averaged within mouse, then averaged across mice. Top, black line depicts mean ΔF/F_0_ Z-score, pink depicts s.e.m. Bottom, heatmap showing signal for individual mice. **E** As a control to test whether vMCs were activated non-specifically by abrupt motor behavior onset, ΔF/F_0_ Z-scores for the same N = 5 mice from (D) during the same behavioral session were aligned to time of offensive attack on an interacting conspecific, averaged within mouse, then averaged across mice. Top, black line depicts mean ΔF/F_0_ Z-score, green depicts s.e.m. Bottom, heatmap showing signal for individual mice. **F** Area under the curve from −2 s to +2 s aligned to either rearing (pink) or attack (green) for mice in (D-E) shows significantly greater vMC activity associated with rearing than attack during the same behavioral social interaction session, suggesting vMCs are not non-selectively activated by any locomotor onset.

### Ventral MCs functionally target dDG granule cells

Electron microscopy has revealed that most vMC axonal synaptic targets in the dDG are on spines of granule cell dendrites in the inner molecular layer (45, 63). To test the functionality of this longitudinal circuit, we used AAV1 as an anterograde transsynaptic tracer of functional synapses, which has been demonstrated in a variety of neuronal circuits (64, 65). We infused AAV1-hSyn-Cre unilaterally into the ventral hilus to serve as the seed region and infused Cre-dependent mCherry (AAV-DIO-mCherry) into either ipsilateral or contralateral dDG to report Cre recombination (**Figure 3A**). AAV-DIO-mCherry infusion did not express mCherry in the absence of ventral hilar AAV1-Cre, suggesting that detectable fluorophore is due to Cre-mediated recombination and not leak expression (**Figure 3B, left**). The addition of AAV1-Cre in either the contralateral (**Figure 3B, middle**) or ipsilateral (**Figure 3B, right**) ventral hilus resulted in mCherry+ granule cells in the dDG, along with some labeling of dorsal hilar neurons. Because this method likely also captured longitudinal projecting vCA3c neurons (66), we used the intersectional targeting technique shown in Figure 1 to express channelrhodopsin selectively in vMCs and performed whole-cell patch-clamp recordings from dDG granule cells while stimulating channelrhodopsin+ terminals in the inner molecular layer using 473 nm blue light at 20 Hz (**Supplementary Figure 2**) (49). Action potentials were readily detected upon depolarization of granule cells membrane potential to approximately −60 mV. These data support functional monosynaptic connectivity between vMCs and dDG granule cells, and are consistent with a recent report from Houser et al. showing functional connectivity in the opposite direction, i.e. from dorsal MCs projecting to ventral granule cells (49).

**Figure 3.**
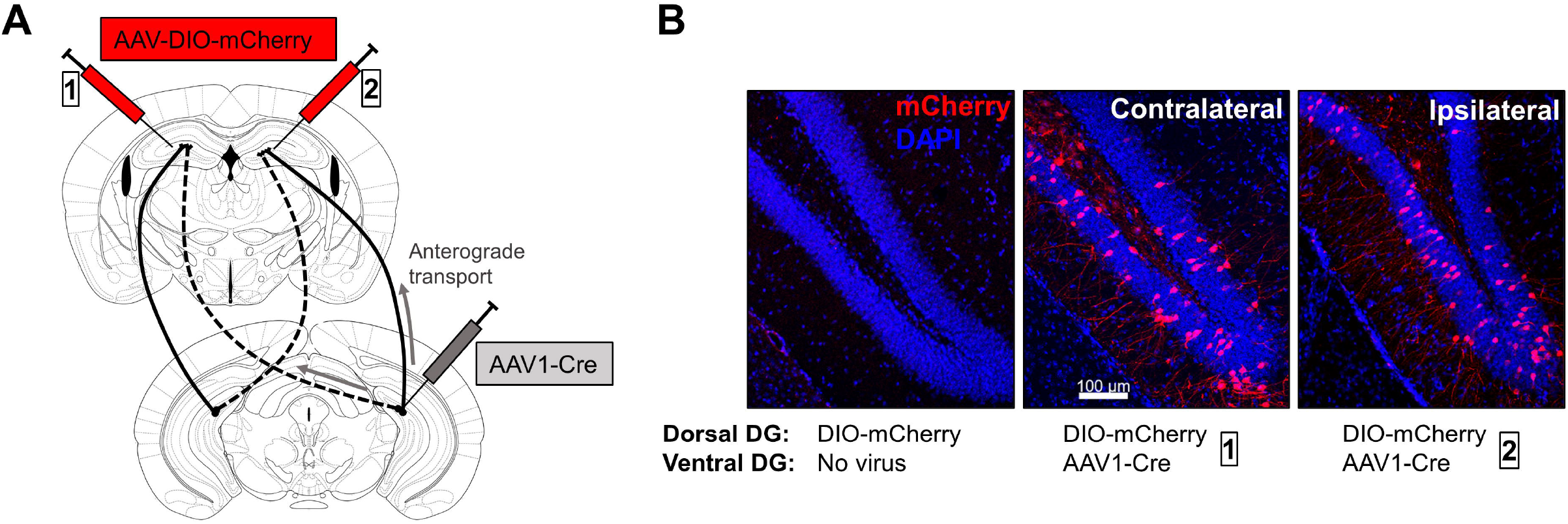
AAV1-mediated anterograde transsynaptic tracing reveals functional synapses between ventral mossy cells and dorsal granule cells. A AAV1-Cre was infused unilaterally into ventral dentate gyrus (DG) hilus, and Cre-dependent mCherry (AAV-DIO-mCherry) was infused either contralaterally (box 1) or ipsilaterally (box 2) to dorsal DG to detect anterograde transsynaptic trafficking. **B** AAV-DIO-mCherry does not express mCherry when infused alone (left), however additional infusion of AAV1-Cre in either contralateral (middle) or ipsilateral (right) DG results in dorsal DG granule cell expression of mCherry, supporting functional synaptic connectivity between the ventral hilus and dorsal DG most consistent with longitudinal mossy cell projections.

### Chemogenetic excitation of vMCs activates dDG granule cells and impairs object location memory encoding

We next aimed to model persistent activation of vMCs and test whether this vMC hyperactivation influences a behavior dependent on the dHPC formation and dDG: spatial learning and memory (30-34, 54). We expressed the G_q_-coupled Designer Receptor Exclusively Activated by Designer Drugs (DREADD) construct hM3Dq-mCherry (37) or control mCherry in bilateral vMCs (**Figure 4A**,**B**). Whole cell recordings from vMCs expressing hM3Dq (n = 4 cells from 3 mice) or mCherry controls (n = 3 cells from 2 mice) showed a significantly greater inward current in hM3Dq-expressing vMCs after bath application of the DREADD-ligand 10 μM CNO compared to control (t(5) = 4.24, p = 0.0068) (**Figure 4C**). To determine an appropriate dose of CNO for *in vivo* behavioral experiments, we systemically injected CNO and stained for cFos expression 90 mins later. We observed low levels of cFos in vMCs expressing mCherry (n = 3 mice) or hM3Dq (n = 4 mice) after vehicle treatment or in mCherry-expressing vMCs (n = 4 mice) after 10 mg/kg CNO treatment (**Figure 4D**). cFos expression was robustly increased in hM3Dq-expressing vMCs (n = 5 mice) by 10 mg/kg CNO treatment, as revealed by a significant treatment x virus interaction in a two-way ANOVA (F(1,12) = 33.38, p < 10^−4^; hM3Dq + CNO vs. all other groups: p < 10^−4^, Sidak post-test). However, hM3Dq + CNO-treated mice also demonstrated dense cFos expression in neighboring granule cells, CA3, and CA1 neurons despite their not expressing hM3Dq, suggestive of overly high levels of activation and possible seizure activity. Indeed, careful review of recorded behavior revealed myoclonic jerks in a subset of mice (data not shown). Reducing CNO to 1 mg/kg still significantly activated vMCs in hM3Dq-expressing MCs (n = 3 mice) compared to mCherry-expressing vMCs (n = 3 mice) (t(2.98) = 5.58, p = 0.012) (**Figure 4E**) but did not hyperactivate neighboring granule cells. Consistent with excitatory connectivity *in vivo* between vMCs and dDG granule cells, 1 mg/kg CNO also significantly increased cFos expression in dDG granule cells (t(3.77)=3.21, p = 0.036) (**Figure 4F**). Based on these studies, we selected 1 mg/kg CNO to examine the effects of vMC hyperactivation in behavioral assays.

**Figure 4.**
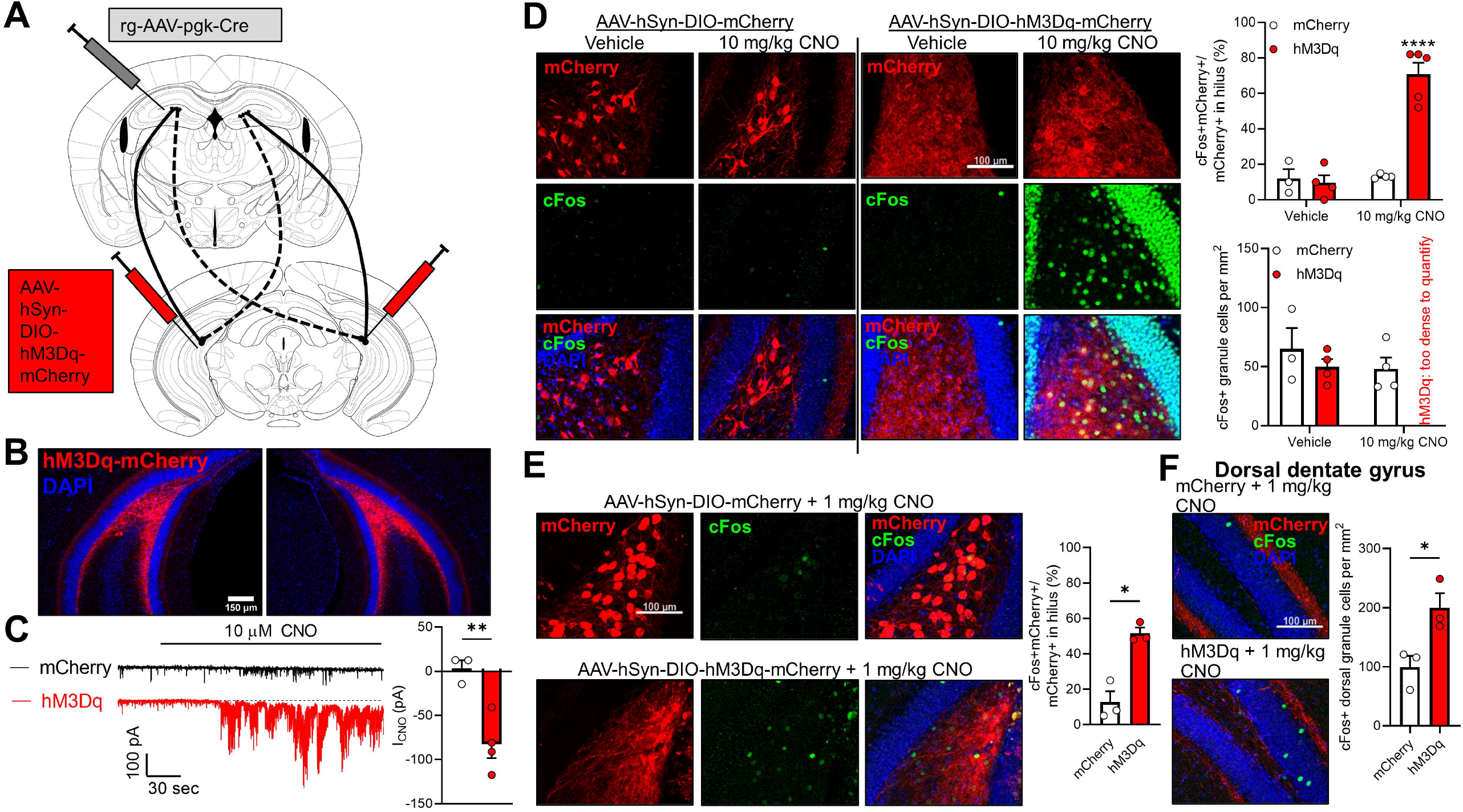
Chemogenetic activation of ventral mossy cells activates dorsal granule cells. A To express hM3Dq-mCherry or mCherry in ventral mossy cells (vMCs), retrograde-AAV-pgk-Cre was unilaterally infused into dorsal dentate gyrus (DG) and AAV-hSyn-DIO-hM3D(Gq)-mCherry or AAV-hSyn-DIO-mCherry was bilaterally infused into ventral DG hilus. **B** Fluorescence microscopy showing representative image of bilateral vMC targeting. **C** Whole cell recordings from vMCs expressing either mCherry (n = 3 cells|2 mice) or hM3Dq-mCherry (n = 4 cells|3 mice) revealed hM3Dq+ neurons showed significantly greater mean inward current after bath application of 10 micromolar clozapine N-oxide (CNO) than control mCherry+ neurons. **p = 0.0068. **D** Mice with vMCs expressing mCherry (n = 3-4) or hM3Dq-mCherry (n = 4-5) were administered vehicle or 10 mg/kg CNO i.p., then perfused 90 mins later and immunostained for cFos. Only vMCs expressing hM3Dq-mCherry strongly expressed cFos, but also strongly expressed cFos in neighboring ventral DG granule cells. ****p < 10^−4^. **E** Same method as in (D) but with reduced dose of CNO (1 mg/kg) showed increase in cFos in vMCs expressing hM3Dq, but no hyperactivation of neighboring granule cells. *p = 0.012. **F** Dorsal DG granule cell cFos was also significantly upregulated in mice with vMC expression of hM3Dq-mCherry treated with 1 mg/kg CNO as compared to mice with vMC expression of mCherry. *p = 0.036.

To test how vMC chemogenetic activation affects spatial learning and memory, we used an object location memory task, which permits temporal separation of encoding and retrieval (42). We focused on encoding because vMC inhibition impaired encoding but not retrieval in previous studies (29). After several days of habituation, mice investigated two identical objects within an arena containing spatial cues (encoding phase), and 24 hrs later returned to the arena where one object was moved to a novel location and the other remained in place. Greater time spent investigating the novel-located object is interpreted as intact spatial memory. Given concerns regarding behavioral tests in outbred mice, we first confirmed that CD-1 mice showed robust object location memory in this paradigm (**Supplementary Figure 3**). We then treated mice expressing vMC hM3Dq or mCherry with 1 mg/kg CNO 30 mins prior to the encoding session. Mice received no treatment prior to the retrieval session 24 hrs later, thus differences during the retrieval session are likely to be due to differences in spatial encoding (**Figure 5A)**. Two-way ANOVA for discrimination index (DI) between the object locations revealed a significant session x virus interaction (n = 10 mCherry mice, n = 9 hM3Dq mice, F(1,17) = 7.64, p = 0.013). Sidak post-hoc tests comparing DI between hM3Dq- and mCherry-expressing mice revealed no significant difference during the encoding session (t(34) = 0.74, p = 0.71) but a significant difference during the retrieval session (t(34) = 2.76, p = 0.018) whereby control mCherry mice spent more time investigating the moved object (**Figure 5B**). The total amount of time spent investigating the objects did not significantly differ between virus groups during encoding (t(15.65) = 0.37, p = 0.72) or retrieval (t(12.65) = 1.40, p = 0.18) sessions (**Figure 5C**). In a 10 min open field test (n = 12 mCherry mice, n = 11 hM3Dq mice) (**Figure 5D**) there were no significant differences in total distance traveled (t(19.74) = 0.27, p = 0.79) or time in the arena center (t(15.13) = 0.68, p = 0.51) between virus groups (**Figure 5E, F**). The total number of rearing events also did not differ between groups (t(20.31) = 0.66, p = 0.52) (**Figure 5g**). Taken together, these data suggest that hyperactivation of vMCs impairs spatial location encoding but does not significantly change exploration of the environment or gross motor output.

**Figure 5.**
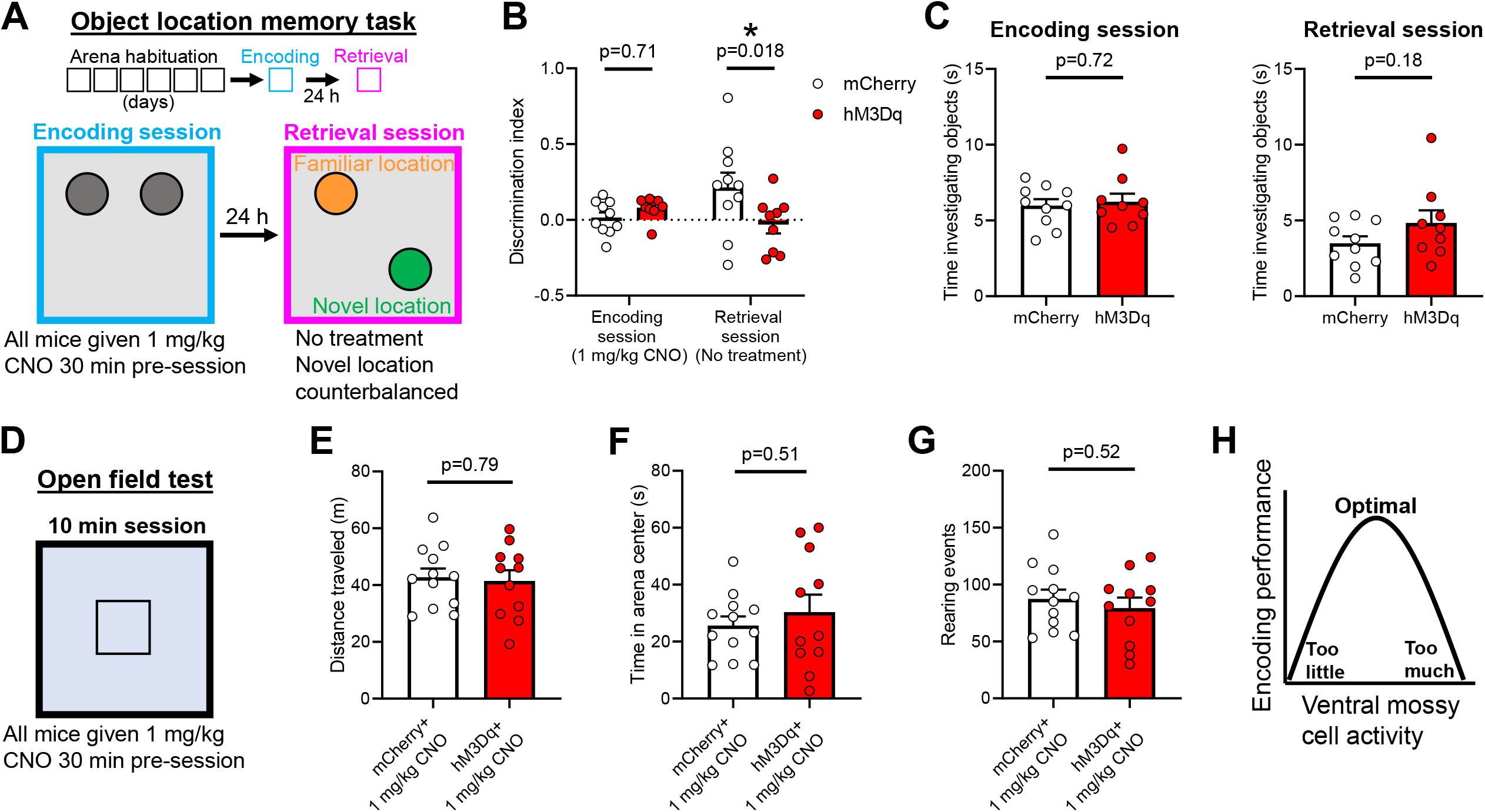
Ventral mossy cell hyperactivation impairs object location memory encoding. A Schematic of the object location memory task. After three days of handling, mice undergo six days of habituation to an arena with spatial cues on the wall. The following day they undergo a 10 min encoding session during which they are exposed to two identical objects. 24 hrs later they are returned to the arena for a 5 min retrieval session where one object has been moved to a novel location and the other remains in the familiar location. 1 mg/kg CNO was administered 30 mins before the encoding session only. **B** CNO administration during encoding reduced object location retrieval in mice with vMC hM3Dq expression (n = 10 mCherry, n = 9 hM3Dq). **C** Total time spent investigating both objects did not significantly differ between the mCherry or hM3Dq-expressing mice during the encoding (left) or retrieval session (right). **D** Mice were later tested in a 10 min open field test in an arena with a novel floor surface 30 mins after 1 mg/kg CNO (n = 12 mCherry, n = 11 hM3Dq). There was no significant difference between mCherry and hM3Dq-expressing mice in total distance travelled (**E**), time spent in arena center (**F**), or number of rearing events (**G**). **H** Schematic suggesting that a specific setpoint of vMC activity exists to promote optimal encoding performance, which is supported by previous work showing that inhibition of vMCs impairs spatial encoding (29) and the current study suggesting that hyperactivation of vMCs also impairs spatial encoding.

## DISCUSSION

Here we used an intersectional viral technique to target, record from, and activate vMCs that densely project to the dDG. Our goal was to interrogate this specific intrahippocampal projection that is anatomically well-suited to distribute localized vHPC (anterior HPC in humans) hyperactivity associated with early psychosis or schizophrenia to disrupt cognitive functions reliant on the dHPC formation and dDG. We found that 1) vMCs in vivo are activated during environmental information gathering, 2) vMCs make functional excitatory synapses with dDG granule cells, 3) chemogenetic activation of vMCs activates dDG granule cells in vivo, and 4) chemogenetic activation of vMCs impairs spatial memory encoding without significantly altering exploratory behaviors. These data suggest that vMC activity associated with rearing is likely related to a role in processing environmental cues but does not directly drive rearing or other locomotor behavior. Next, given the relative importance of the dHPC formation in spatial learning and memory compared to the vHPC (31), our data support the idea that sustained activation of vMCs impairs spatial encoding by aberrantly activating dDG granule cells. Importantly, these results do not preclude the possibility of spatial encoding deficits via non-dDG mechanisms, and future studies will be necessary to target only the vMC to dDG projections Since MCs target both granule cells and inhibitory interneurons, considerable work has addressed whether the net effect of MC activation is to excite or inhibit DG granule cells (23, 24, 29, 52, 67, 68), primarily in the context of seizure propagation. Less attention has been devoted to the role of MCs, especially vMCs, in behavior relevant to neuropsychiatric disease including learning, memory, novelty, and anxiety, however there is growing research interest in these topics. Distinctions between vMCs and dMCs are supported by physiological (69-72), biochemical (27, 28), and anatomical (49, 72, 73) evidence. It thus may not be sufficient to extrapolate findings from dMCs to all MCs across the HPC longitudinal axis. Of note, an elegant recent study using in vivo microscopy showed increased vMC activity in novel versus familiar environments that was not due to locomotion differences (72), in line with our fiber photometry findings.

Our results, taken together with studies showing inhibition of vMCs also impairs encoding in a similar object location memory task (29), supports an overall model that bidirectional perturbation of vMC activity, either too much or too little, degrades spatial encoding (**Figure 5h**), without influencing exploratory or locomotor behavior itself. The point of convergence is likely dDG granule cells, as previous work showed that activation *or* inhibition of dDG but not vDG granule cells impaired contextual encoding (48). Two recent studies illustrate that the effects of vMCs on learning and memory are likely task specific. Fredes *et al*. showed that vMC activation prior to shock-context pairing *enhanced* fear conditioning when the conditioning occurred in a familiar context (72), an important finding because contextual fear conditioning is markedly stronger in novel versus familiar contexts. Another study reported that vMC chemogenetic activation did not significantly impair encoding in an object location memory task in which the delay between encoding and retrieval sessions was only one hr (74) as opposed to the 24 hr delay used in the present study and that of Bui *et al*. (29).

The present results, along with post-mortem evidence for cellular and biochemical changes in human hippocampal area CA4 (75-77) support future investigation into vMC function in rodent models for ventral hippocampal hyperactivation and cognitive deficits seen in schizophrenia. A key goal is to determine why vMC inhibition or excitation impairs spatial encoding. One plausible explanation based on the neuroanatomy of vMC projections to the dDG is through interactions with the medial and/or lateral perforant path from the entorhinal cortex, which terminates in the middle or outer molecular layer on granule cell dendrites, respectively, distal to vMC projection terminals (78, 79) (49). Hypo-or hyperactivation of vMC projections to dorsal granule cells may impair the dDG’s critical function of transforming inputs into sparse ensembles of activated granule cells for pattern separation (80-85). Future studies using mouse models with genetic lesions conferring high risk for schizophrenia in humans that show vHPC hyperactivity and object location memory deficits, such as 22q11.2 deletion models (86, 87), are well-suited to test whether modulating vMC longitudinal projections may rescue spatial memory deficits. Finally, chemogenetic manipulations in this study were acute, and it has been shown that learning and memory changes following viral MC ablations are surprisingly transient (52). Given aberrant neurodevelopmental processes in schizophrenia, it will be important to test whether chronic vMC inhibition or excitation impairs spatial memory in a similar manner to acute manipulations.

## ACKNOWLEDGEMENTS

This work was supported by National Institutes of Health grants MH116339 (A.S.L.) and NS107424 (C.Z.), and the Vanderbilt Department of Psychiatry and Behavioral Sciences. Experiments and data analysis were performed in part using the Vanderbilt University Medical Center Cell Imaging Shared Resource (supported by NIH grants CA68485, DK20593, DK58404, DK59637 and EY08126). We thank Marina Picciotto, Ph.D. and Ye Han, Ph.D. for helpful comments on this work. This work has been posted as a preprint to bioRxiv.

