## Supplemental Information for "Modeling intrahippocampal effects of anterior hippocampal hyperactivity relevant to schizophrenia using chemogenetic excitation of long axis-projecting mossy cells in the mouse dentate gyrus"

**SUPPLEMENTAL MATERIALS AND METHODS**

**Adeno-associated viruses**

The following adeno-associated virus particles (AAVs) were purchased from Addgene (Watertown, MA). Citations for published constructs are provided in the main manuscript.

1. AAV.pgk.Cre was a gift from Patrick Aebischer (Addgene viral prep # 24593-AAVrg; http://n2t.net/addgene:24593; RRID:Addgene_24593). Viral titer = 9.3 x 10^12^ genome copies (GC)/mL.

2. pAAV.Syn.Flex.GCaMP6f.WPRE.SV40 was a gift from Douglas Kim & GENIE Project (Addgene viral prep # 100833-AAV1; http://n2t.net/addgene:100833; RRID:Addgene_100833). Viral titer = 1.9 x 10^13^ GC/mL.

3. pAAV-hSyn-DIO-mCherry was a gift from Bryan Roth (Addgene viral prep # 50459-AAV8; http://n2t.net/addgene:50459; RRID:Addgene_50459). Viral titer = 2.6 x 10^13^ GC/mL.

4. pAAV-hSyn-DIO-hM3D(Gq)-mCherry was a gift from Bryan Roth (Addgene viral prep # 44361-AAV8; http://n2t.net/addgene:44361; RRID:Addgene_44361). Viral titer = 2.0 x 10^13^ GC/mL.
5. pENN.AAV.hSyn.Cre.WPRE.hGH was a gift from James M. Wilson (Addgene viral prep # 105553-AAV1; http://n2t.net/addgene:105553 ; RRID:Addgene_105553). Viral titer = 1.9 x 10^13^ GC/mL.

6. pAAV-EF1a-double floxed-hChR2(H134R)-mCherry-WPRE-HGHpA was a gift from Karl Deisseroth (Addgene viral prep # 20297-AAV8; http://n2t.net/addgene:20297 ; RRID:Addgene_20297). Viral titer = 1.9 x 10^13^ GC/mL.

**Stereotaxic surgery and viral infusion**

Male CD-1 mice were anesthetized using inhaled isoflurane (Piramal Critical Care, Telangana, India) administered by a tabletop anesthesia machine (VetEquip, Livermore, CA) with thermal support provided by Deltaphase isothermal pads (Braintree Scientific, Braintree, MA) and mounted in a stereotaxic frame (Kopf Instruments, Tujunga, CA). Adeno-associated virus (0.2 – 0.4 μL, depending on target region) was infused at a rate of 50 nL/min using a 2 μL syringe (Hamilton, Reno, NV) to the following structures:

1) Dorsal dentate gyrus (DG) inner molecular layer: Anterior/posterior (AP): -1.94 mm, medial/lateral (ML): 1.20 mm left; dorsal/ventral (DV): -2.50 mm.

2) Ventral DG hilus: AP: -3.40 mm, ML: ±3.00 mm, DV: -3.50 mm.

The syringe was left in place for an additional 3 min to reduce backflow. Fiber photometry cannulas were targeted to the same coordinates as ventral DG hilus and cemented in place with C&B-Metabond (Parkell, Edgewood, NY). The skin was closed with VetBond (3M, Saint Paul, MN). Mice were housed for at least three weeks prior to experimentation to allow for viral expression. Targeting was confirmed by fluorescence microscopy. Mice with either unilateral or bilateral ventral mossy cell expression were included in behavioral analyses due to extensive bilateral projections.

**Fluorescent immunostaining, microscopy, and cell counting**

Mice were terminally anesthetized with pentobarbital sodium (Vortech Pharmaceuticals, Dearborn, MI) then perfused intracardially with ice-cold phosphate buffered saline (PBS, 0.1 M, pH 7.3) followed by ice-cold 4% paraformaldehyde (PFA) in PBS. Brains were removed and postﬁxed for 48 h in 4% PFA at 4 ℃, then stored in PBS until sectioning. 40 μm sections were cut on a vibrating microtome (Leica, Buffalo Grove, IL). For immunostaining, tissue sections were permeabilized and blocked in 0.3% Triton X-100 solution and 3% normal donkey serum (NDS, Jackson ImmunoResearch, West Grove, PA) in PBS (blocking buffer) for 2 hours at room temperature followed by incubation in primary antibody. Primary antibodies used were mouse anti-calretinin (MAB1568, Millipore, Burlington, MA, diluted in blocking buffer) and rabbit anti-cFos (226 003, Synaptic Systems, Goettingen, Germany, diluted in 0.1% Triton X-100 and 1% NDS in PBS), both diluted 1:1000 with overnight incubation at 4 ℃. Sections were rinsed in PBS, then incubated in secondary antibody (Alexa Fluor 488 Donkey Anti-Mouse, Alexa Fluor 488 Donkey Anti-Rabbit, or Alexa Fluor 647 Donkey Anti-Rabbit) diluted 1:1000 in blocking buffer for 2 hours at room temperature. Sections were rinsed in PBS, then incubated in 4′,6-diamidino-2-phenylindole (DAPI, Millipore) diluted 1:5000 in PBS at room temperature for 5 mins. Sections were again washed in PBS, then mounted on slides using Fluoromount G (Electron Microscopy Sciences, Hatfield, PA). Widefield images were acquired with an AF6000 LX fluorescent microscopy system (Leica, Buffalo Grove, IL) equipped with an HCX PL FLUOTAR 10.0x objective (NA = 0.30), while confocal images were acquired with an LSM 710 META Inverted microscope (Zeiss, White Plains, NY) equipped with a 20x Plan-Apochromat objective (NA = 0.8). Acquisition software was Leica LAS AF for widefield images and Zeiss Zen for confocal images. To quantify the overlap between GCaMP6f and calretinin, two neighboring coronal sections were imaged as Z stacks and collapsed using maximum projection. The number of hilar GCaMP6f+, calretinin+, and GCaMP6f+calretinin+ double labeled neurons were counted, enabling calculation of GCaMP6f+calretinin+/GCaMP6f+ and GCaMP6f+calretinin+/calretinin+. These values were averaged across two sections to obtain a single value for each mouse, and these single values were averaged across mice. For off-target GCaMP6f quantification, the same approach was used and the number of GCaMP6f+ neurons was divided by the area of the dentate gyrus granule cell layer as delineated by DAPI staining or area CA3. Quantification of hilar cFos+mCherry+/mCherry+ neurons to calculate fraction of hilar cells activated by DREADD strategy was performed in a similar fashion, as was quantification of cFos+ neurons per unit area in dentate gyrus granule cells of the ventral hippocampus. To quantify cFos+ neurons in the dorsal hippocampus, the same technique was used except cFos+ neurons were counted in bilateral hippocampus for each slice and normalized to granule cell layer area, averaged to get a single value per slice, and then averaged with a second slice so that each mouse contributed a single data point consisting typically of 4 hippocampi.

**Electrophysiology**

*Validation of DREADD activation*

Mice were rapidly decapitated under deep isoflurane anesthesia and 300-µm horizontal sections containing the ventral DG were prepared in *N*-methyl-D-glucamine cutting solution (in mM): 93 *N*-methyl-D-glucamine, 20 HEPES, 2.5 KCl, 0.5 CaCl_2_, 10 MgCl_2_, 1.2 NaH_2_PO_4_, 25 glucose, 5 Na-ascorbate, and 3 Na-pyruvate. Slices recovered in 30° cutting solution for 10 minutes and then in 23° artificial cerebrospinal fluid for at least 60 minutes prior to recording. All solutions were continuously bubbled with 95/5% O_2_/CO_2_. Slices were transferred to a 30° recording chamber perfused at 2 mL/min with artificial cerebrospinal fluid (in mM): 119 NaCl, 2.5 KCl, 2.5 CaCl_2_, 1.3 MgCl2, 1 NaH_2_PO_4_, 11 glucose, and 26 NaHCO_3_. Mossy cells were identified within the hilus, expressed red fluorescent protein, and were patched with borosilicate glass pipettes (4-6 MΩ). Whole cell access was obtained, and cells were dialyzed for 5 minutes with internal solution (in mM): 125 K-gluconate, 4 NaCl, 10 HEPES, 4 MgATP, 0.3 NaGTP, 10 Tris-phosphocreatine. Mossy cells were voltage-clamped at -75 mV and 10 µM CNO was perfused in the bath for 5 minutes. Recordings were acquired with a Multiclamp 700B amplifier (Molecular Devices, Sunnyvale, CA), filtered at 2 kHz and digitized at 10 kHz. The mean of the holding current was quantified before and after CNO perfusion using pClamp 10.4 software (Axon Instruments, Union City, CA).

*Optical activation of ventral mossy cell terminals in dorsal dentate gyrus*

Brain slices were prepared as previously described (1, 2). Mice were deeply anesthetized and transcardially perfused with an N-methyl-D-glucamine (NMDG)-based ice-cold solution (components below) and decapitated. Coronal brain slices (300-320 µm thickness) containing dorsal hippocampus were prepared with a vibratome (Leica VT 1200S, Leica Biosystems Inc) in an NMDG dissection solution (mM: NMDG 92, KCl 2.5, CaCl_2_ 0.5, NaH_2_PO_4_ 1.25, HEPES 20, MgSO_4_ 10, NaHCO_3_ 30, glucose 25, Thiourea 2, Na-ascorbate 5, Na-pyruvate 3, pH 7.3-7.4 and O_2_ 95%+5%CO_2_ bubbled) (3) and slices were later incubated in a chamber at 35-36°C for 40 min with continuously oxygenated ACSF (see below for components). Slices remained at room temperature for at least 1 hour before electrophysiological recordings at room temperature. Whole-cell patch-clamp recordings were made from dorsal hippocampal dentate gyrus granule neurons by using a Nikon infrared/DIC microscope (Eclipse FN1, Nikon Corp. Inc., Melville NY), and slices were continuously superfused (flow speed 1-1.5 ml/min) with an ACSF solution (containing [in mM]: 126 NaCl, 2.5 KCl, 1.25 NaH_2_PO_4_, 2 CaCl_2_, 2 MgCl_2_, 26 NaHCO_3_, 10 D-glucose, pH 7.4) bubbled with 95% O_2_/5% CO_2_. Filled electrodes had resistances of 2~5 MΩ with one internal solution (consisted of [in mM]: 120 K-gluconate, 11 KCl, 1 MgCl_2_, 1 CaCl_2_, 0.6 EGTA, 10 HEPES, 2 Na-ATP, 0.6 Na-GTP, 10 K-creatine-phosphate, pH 7.3 (4, 5)) for spontaneous (s) and α-amino-3-hydroxy-5-methyl-4-isoxazolepropionic acid receptor (AMPAR)-mediated sEPSCs, recorded at a holding potential -55.8mV (Cl^-^ reversal potential). To record action potentials (APs), current-clamp mode was used with the same internal solution for sEPSC recordings. Access resistance (Ra, voltage-clamp mode) was continuously monitored during recordings and recordings with Ra larger than 25 MΩ or 20% change were discarded. The blue laser light (5 ms pulse duration, 20 Hz, 1 s duration) was delivered through a fiberoptic cable to dentate gyrus within brain slices for stimulating axon terminals containing ChR2 within dentate gyrus, controlled by a DPSS laser (MBL-III-473 (100mW, Ultralazers Co., Inc)) and the timing of laser delivery was controlled by Clampex 10 software (Molecular Devices Inc., Union City, CA). Data were collected using one multiClamp 700B amplifier and Clampex 10 software (Molecular Devices Inc., Union City, CA) and filtered at 2 kHz, and digitized at 20 kHz using a Digidata 1440A (Molecular Devices Inc., Union City, CA).

**Fiber photometry**

Fiber photometry cannula (Doric Lenses, Quebec, Canada) implanted male CD-1 mice expressing GCaMP6f in vMCs underwent *in vivo* calcium recording during resident-intruder interactions described above. Photometry was performed using the Doric Lenses system controlled by Doric Neuroscience Studio (DNS) version 5.3, consisting of 405 nm and 465 nm light-emitting diodes (LEDs) run by an LED driver and routed through a 4-port fluorescent minicube. GCaMP6f output excited by 405 nm (calcium-independent signal) and 465 nm (calcium-dependent signal) LEDs were measured by a Newport Visible Femtowatt Photoreceiver and separated using lock-in demodulation. Behavior was time-locked with photometry signal by USB 3.0 color camera (Doric Lenses) input to the photometry console. Signal processing was performed by DNS photometry analyzer. ΔF/F_0_ was calculated for the 405 nm and 465 nm channels independently using a least mean square fit of the whole trace, and then ΔF/F_0_ (405 nm) subtracted from ΔF/F_0_ (465 nm) to yield a corrected trace. This trace was subsequently lowpass filtered at 2 Hz (6) to yield the final bulk calcium signal and Z-normalized across the entire social interaction. Videos were annotated manually to mark time of non-social exploratory rearing (centered around rearing apogee) and time of offensive attack. We only included attacks occurring in isolation (single attack) or the first in a series of multi-attack epochs to avoid contamination from subsequent attacks. ΔF/F_0_ Z-scores from -2 s to +2s for each rear or attack were averaged for each mouse, then averaged across mice. Data analysis was performed using custom written scripts in Matlab R2019a (MathWorks, Natick, MA). Only mice with histologically verified cannula and GCaMP6f targeting were analyzed.

**SUPPLEMENTAL FIGURES AND LEGENDS**

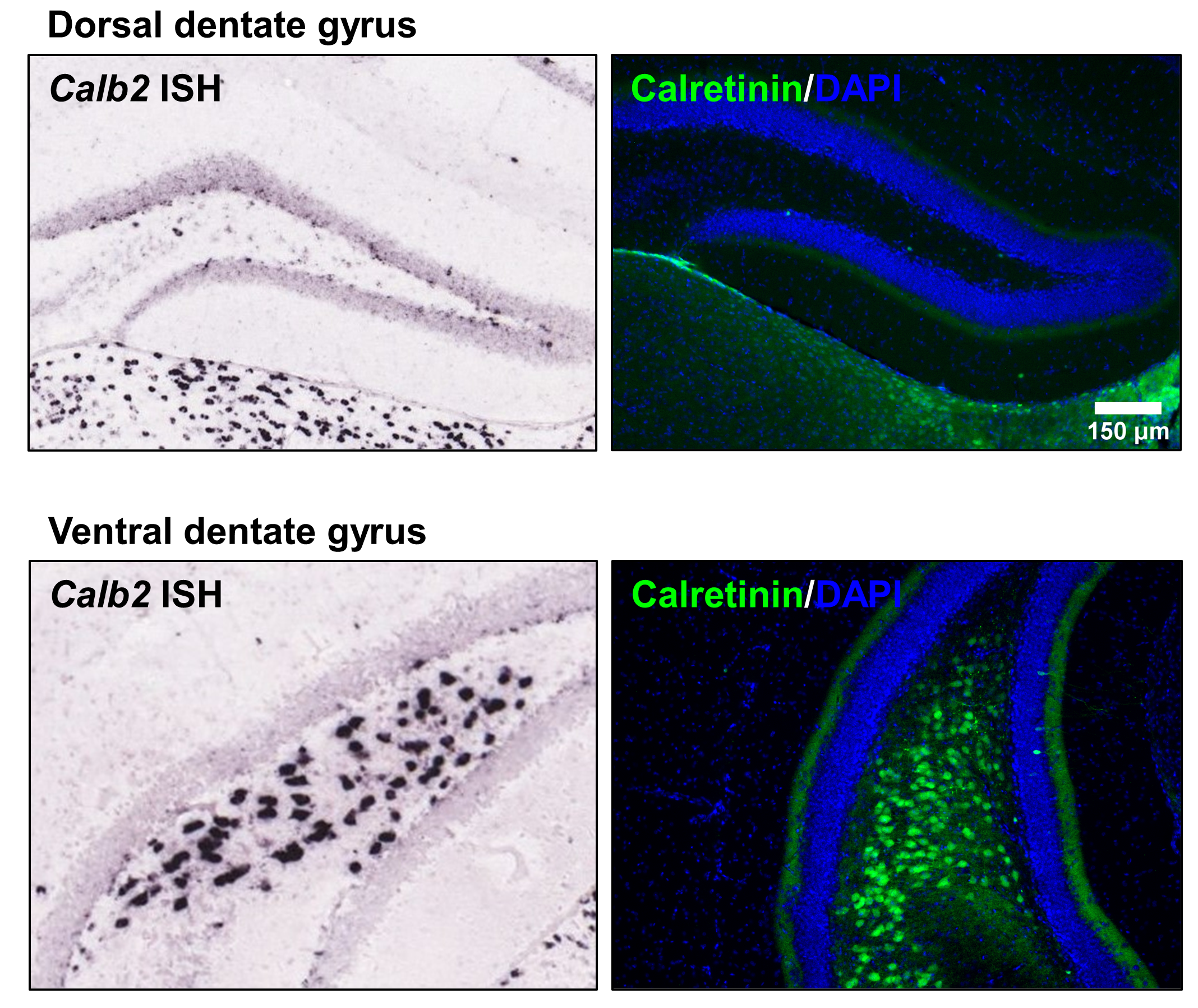

**Supplemental Figure 1.** Calretinin expression in the dorsal and ventral hippocampal hilus. In situ hybridization (ISH) for the gene encoding calretinin, *Calb2* (left), and immunofluorescent images for calretinin protein (right) confirmed previous studies that ventral mossy cells (bottom) but not dorsal mossy cells (top) express calretinin mRNA and protein. In situ hybridization is from the Allen Mouse Brain Atlas (7, 8).

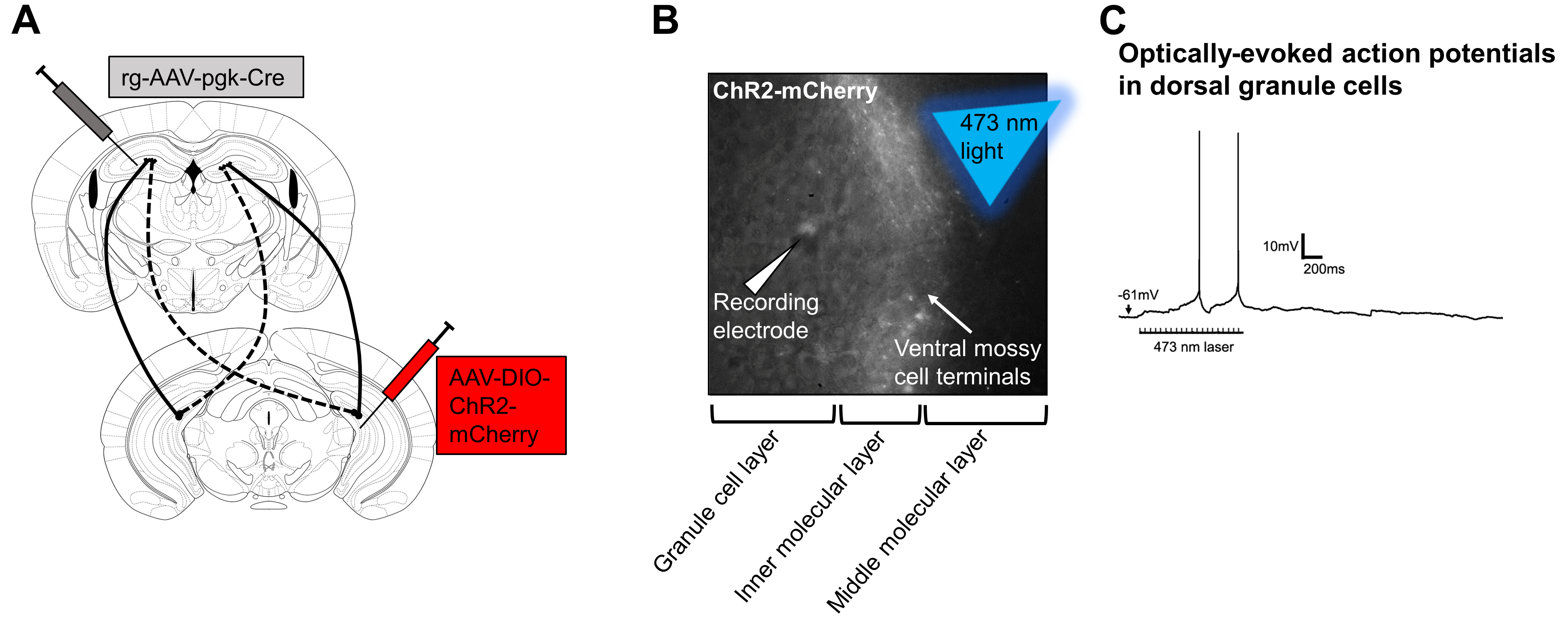

**Supplemental Figure 2.** Optogenetic excitation of ventral mossy cells terminals in the dorsal dentate gyrus elicits action potentials in dorsal dentate gyrus granule cells. **A** Channelrhodopsin-mCherry (AAV-EF1a-double floxed-hChR2(H134R)-mCherry-WPRE-HGHpA) was expressed in ventral mossy cells projecting to the contralateral dorsal dentate gyrus using an intersectional targeting technique. **B** In slices, whole-cell patch-clamp recordings from dorsal dentate gyrus granule cells were performed while stimulating channelrhodopsin-mCherry-expressing ventral mossy cell terminals in the dorsal dentate gyrus inner molecular layer using a blue laser (473 nm, 5 ms pulse duration, 20 Hz for 1 s). **C** Action potentials were readily elicited from dorsal dentate gyrus granule cells (representative image, N = 3 cells | 2 mice).

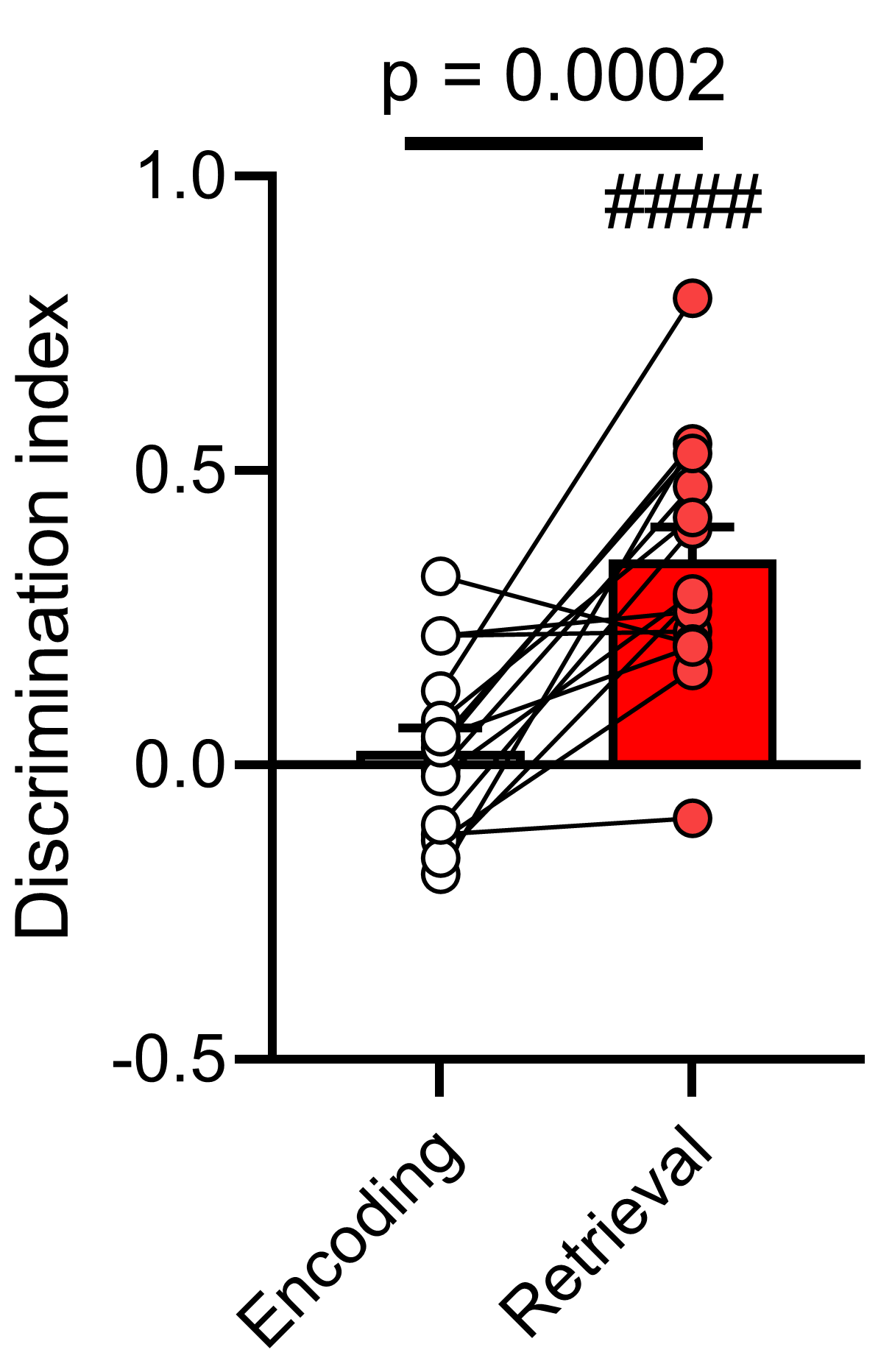

**Supplemental Figure 3.** CD-1 wildtype mice show intact object location memory 24 hrs after encoding as evidenced by a significant increase in discrimination index (DI) between encoding and retrieval sessions (N = 14, t(14) = 4.94, p = 0.0002). ####p < 10^-4^ by one-sample t test versus DI = 0 (t(14) = 6.37).
